## Supplemental Tables and Figures for "Trehalose increases tomato drought tolerance, induces defenses, and increases resistance to bacterial wilt disease"

**Supplemental Table 1. Strains and primers used in this study**

| ***Strain*** | ***Notes*** | ***Reference*** |
|---|---|---|
| Wild type GMI1000 | *Ralstonia solanacearum* phylotype one sequevar 18, isolated from tomato in French Guyana | (Boucher, Barberis et al. 1985) |
| GMI1000 Δ*treA* | GMI1000 trehalase mutant cannot degrade trehalose; Spectinomycin/streptomycin-resistant | (MacIntyre, Barth et al. 2019) |
| ***Primer Name*** | ***Sequence*** | ***Reference*** |
| loxA F | TGGTAGACCACCAACACGAA | (Milling, Babujee et al. 2011) |
| loxA R | GACCAAAACGCTCGTCTCTC | " |
| osmF | TGTACCACGTTTGGAGGACA | " |
| osmR | ACCAGGGCAAGTAAATGTGC | " |
| PR1b B F | TTGGTGACTGCGGGATGA | " |
| PR1b B R | GGCGGCGGCTAGGTT T | " |
| PR1a B F | GAGGGCAGCCGTGCAA | " |
| PR1a B R | CACATTTTTCCACCAACACATTG | " |
| GluA F | TCA GCA GGG TTG CAA AAT CA | " |
| GluA R | CTCTAGGTGGGTAGGTGTTGGTTAA | " |
| Pin2 F | TGATGCCAAGGCTTGTACTAGAGA | " |
| Pin2 R | AGCGGACTTCCTTCTGAACGT | " |
| ACO5 F | AGATGGGCATTGGGTGAACA | " |
| ACO5 R | TTCAGCCATCACTCGGTGTC | " |
| DnaJ F | ATGAAGCGCCAGATACCATC | " |
| DnaJ R | TCAAGGCTCAATGTGTGCTC | " |
| Actin F | TCAGCAACTGGGATGATATG | " |
| Actin R | TTAGGGTTGAGAGGTGCTTC | " |
| rd22_F | ACGTGGCGTTATTTTTCCTG | (Yamaguchi-Shinozaki and Shinozaki 1993) |
| rd22_R | ATCTCCGGCATCTTCTCTGA | " |
| dhn_tas_F | CACCATGAGGGGCAACAGCA | (Kissoudis, Seifi et al. 2016) |
| dhn_tas_R | TCACCTTCATGTTGTCCAGGCATC | " |

**Supplemental Table 2. Concentration of hormones, oxylipins, and phenylpropanoids in tomato xylem sap following various treatments**

| Compound (nM)[a] | Healthy Avg[b] | WT avg[c] | WT FC[d] | WT p-val[e] | 6 hr tre avg[f] | 6 hr tre FC | 6 hr tre p-val | 24 hr tre avg[g] | 24 hr tre FC | 24 hr tre p-val | 48 hr tre avg[h] | 48 hr tre FC | 48 hr tre p-val |
|---|---|---|---|---|---|---|---|---|---|---|---|---|---|
| 12,13-diHOM | 0.83 | 1.30 | 2 | 0.04 | 1.26 | 2 | 0.27 | 2.69 | 3 | 0.00 | 1.50 | 2 | 0.02 |
| 12,13-Ep-9KOM | 5.38 | 6.86 | 1 | 0.24 | 7.93 | 1 | 0.02 | 4.82 | 1 | 0.64 | 7.62 | 1 | 0.04 |
| 12OH-JA | 1.43 | 1.70 | 1 | 0.63 | 5.21 | 4 | 0.30 | 1.38 | 1 | 0.84 | 1.17 | 1 | 0.06 |
| 12OH-JA-Ile | 12.37 | 15.23 | 1 | 0.81 | 20.67 | 2 | 0.29 | 22.97 | 2 | 0.14 | 57.67 | 5 | 0.00 |
| 12-OPDA | 35.86 | 47.89 | 1 | 0.44 | 50.75 | 1 | 0.43 | 138.72 | 4 | 0.00 | 91.84 | 3 | 0.01 |
| 13-HOD | 4.66 | 8.52 | 2 | 0.00 | 5.09 | 1 | 0.66 | 9.02 | 2 | 0.01 | 5.49 | 1 | 0.14 |
| 13-KOD | 4.50 | 10.78 | 2 | 0.01 | 6.27 | 1 | 0.31 | 9.72 | 2 | 0.00 | 5.81 | 1 | 0.12 |
| 13,12-KODA | 0.27 | 0.75 | 3 | 0.34 | 0.11 | 0 | 0.37 | 4.89 | 18 | 0.00 | 0.22 | 1 | 0.80 |
| 13,12-KOMA | 1.78 | 3.18 | 2 | 0.23 | 1.23 | 1 | 0.47 | 9.67 | 5 | 0.00 | 2.23 | 1 | 0.58 |
| 2-HOD | 0.24 | 0.35 | 1 | 0.44 | 0.36 | 1 | 0.18 | 0.30 | 1 | 0.55 | 0.29 | 1 | 0.66 |
| 9,10,11-THOD | 88.94 | 191.22 | 2 | 0.02 | 134.71 | 2 | 0.09 | 214.49 | 2 | 0.00 | 153.30 | 2 | 0.02 |
| 9,10,11-THOM | 237.57 | 811.16 | 3 | 0.00 | 430.01 | 2 | 0.02 | 572.57 | 2 | 0.00 | 379.05 | 2 | 0.00 |
| 9,10,13-THOD | 3.47 | 4.47 | 1 | 0.28 | 9.91 | 3 | 0.00 | 5.78 | 2 | 0.00 | 6.06 | 2 | 0.02 |
| 9,10,13-THOM | 62.08 | 100.56 | 2 | 0.03 | 147.39 | 2 | 0.00 | 128.47 | 2 | 0.00 | 96.87 | 2 | 0.01 |
| 9,10-diHOM | 0.95 | 1.25 | 1 | 0.43 | 1.48 | 2 | 0.52 | 3.64 | 4 | 0.00 | 1.02 | 1 | 0.86 |
| 9,12,13-THOD | 39.77 | 50.62 | 1 | 0.18 | 43.68 | 1 | 0.56 | 84.76 | 2 | 0.00 | 60.09 | 2 | 0.02 |
| 9,12,13-THOM | 127.69 | 231.37 | 2 | 0.01 | 153.78 | 1 | 0.36 | 305.89 | 2 | 0.00 | 179.49 | 1 | 0.04 |
| 9-HOD | 6.15 | 21.29 | 3 | 0.01 | 10.27 | 2 | 0.02 | 14.60 | 2 | 0.00 | 8.64 | 1 | 0.17 |
| 9-HOT | 3.04 | 9.78 | 3 | 0.03 | 5.02 | 2 | 0.16 | 9.75 | 3 | 0.00 | 4.87 | 2 | 0.12 |
| 9-KOD | 4.35 | 8.90 | 2 | 0.15 | 6.60 | 2 | 0.34 | 19.32 | 4 | 0.00 | 12.16 | 3 | 0.00 |
| 9-KOT | 1.06 | 3.68 | 3 | 0.17 | 5.51 | 5 | 0.05 | 7.61 | 7 | 0.00 | 3.90 | 4 | 0.05 |
| 9OH-10KOM | 8.81 | 5.36 | 1 | 0.07 | 10.81 | 1 | 0.15 | 9.05 | 1 | 0.88 | 8.97 | 1 | 0.92 |
| 9OH-12KOD | 0.20 | 0.13 | 1 | 0.51 | 0.27 | 1 | 0.53 | 0.69 | 4 | 0.00 | 0.34 | 2 | 0.02 |
| 9OH-12KOM | 1.88 | 2.48 | 1 | 0.03 | 2.96 | 2 | 0.24 | 3.97 | 2 | 0.00 | 2.11 | 1 | 0.36 |
| 9-oxo-NA | 13.94 | 17.47 | 1 | 0.37 | 16.53 | 1 | 0.56 | 16.06 | 1 | 0.51 | 17.25 | 1 | 0.32 |
| ABA | 8.20 | 49.59 | 6 | 0.03 | 5.71 | 1 | 0.60 | 40.37 | 5 | 0.00 | 9.72 | 1 | 0.73 |

| Compound (nM)[a] | Healthy Avg[b] | WT avg[c] | WT FC[d] | WT p-val[e] | 6 hr tre avg[f] | 6 hr tre FC | 6 hr tre p-val | 24 hr tre avg[g] | 24 hr tre FC | 24 hr tre p-val | 48 hr tre avg[h] | 48 hr tre FC | 48 hr tre p-val |
|---|---|---|---|---|---|---|---|---|---|---|---|---|---|
| AZA | 14.08 | 19.39 | 1 | 0.11 | 19.21 | 1 | 0.00 | 20.44 | 1 | 0.00 | 18.03 | 1 | 0.01 |
| BA | 7.18 | 12.37 | 2 | 0.13 | 11.34 | 2 | 0.19 | 13.29 | 2 | 0.00 | 9.74 | 1 | 0.02 |
| CA | 7.49 | 5.19 | 1 | 0.52 | 5.30 | 1 | 0.56 | 22.69 | 3 | 0.00 | 14.19 | 2 | 0.16 |
| COUMA | 2.04 | 149.10 | 73 | 0.05 | 2.42 | 1 | 0.83 | 5.22 | 3 | 0.11 | 0.00 | 0 | 0.12 |
| dnOPDA | 0.12 | 0.21 | 2 | 0.25 | 0.26 | 2 | 0.02 | 0.53 | 4 | 0.00 | 0.44 | 4 | 0.00 |
| IAA | 3.26 | 36.76 | 11 | 0.03 | 2.84 | 1 | 0.59 | 4.01 | 1 | 0.29 | 2.20 | 1 | 0.09 |
| JA | 16.71 | 9.42 | 1 | 0.22 | 26.64 | 2 | 0.33 | 49.41 | 3 | 0.01 | 71.36 | 4 | 0.00 |
| JA-Ile | 26.59 | 3.67 | 0 | 0.09 | 35.48 | 1 | 0.62 | 60.08 | 2 | 0.04 | 103.68 | 4 | 0.00 |
| OPC-4 | 0.31 | 0.53 | 2 | 0.09 | 0.52 | 2 | 0.13 | 1.09 | 4 | 0.00 | 1.05 | 3 | 0.00 |
| p12,13-DiHOD* | 0.16 | 0.91 | 6 | 0.00 | 0.85 | 5 | 0.05 | 0.82 | 5 | 0.00 | 0.18 | 1 | 0.69 |
| p9,10-DiHOD* | 0.63 | 1.75 | 3 | 0.02 | 1.48 | 2 | 0.04 | 1.73 | 3 | 0.01 | 0.91 | 1 | 0.42 |
| p9OH-TAN* | 3.62 | 3.35 | 1 | 0.62 | 3.36 | 1 | 0.71 | 8.86 | 2 | 0.00 | 4.07 | 1 | 0.46 |
| pOTD* | 0.47 | 0.42 | 1 | 0.72 | 0.52 | 1 | 0.71 | 1.19 | 3 | 0.00 | 0.36 | 1 | 0.33 |
| SA | 67.53 | 31.73 | 0 | 0.21 | 55.50 | 1 | 0.69 | 173.44 | 3 | 0.01 | 55.09 | 1 | 0.65 |
| TA | 6.03 | 10.56 | 2 | 0.04 | 7.94 | 1 | 0.19 | 17.41 | 3 | 0.00 | 7.35 | 1 | 0.23 |
| TAN | 0.54 | 0.77 | 1 | 0.32 | 0.53 | 1 | 0.91 | 0.74 | 1 | 0.01 | 0.60 | 1 | 0.51 |

[a] Compound color key: Blue fill= 13-LOX pathway derived compound; Red fill= 9-LOX pathway derived compound; Purple fill= phenylpropanoid pathway derived compound; Gray fill= other; metabolites were measured with LC/MS/MS

[b] Xylem sap harvested from mock-inoculated/treated healthy BB plants

[c] Sap harvested from DI=1 *Rs* infected plants

[d] Fold-change values ≥ 2 are shaded in light red with dark red text.

[e] P-value derived from t-test of treatment compared to healthy; *P*<0.5 was shaded yellow with darker yellow text.

[f-g] Xylem sap harvested from uninoculated, trehalose treated plants 6, 24, and 48 h post-treatment

**Supplemental Table 3. Metabolite key for hormone analysis**

| Q1[a] | Q3[b] | RT[c] | Metabolite | Formal Name |
|---|---|---|---|---|
| 313.3 | 183.3 | 10.16 | 12,13-DiHOM | threo-12,13-dihydroxy-9(*Z*)-octadecenoic acid |
| 309.4 | 208.9 | 12.46 | (*E*)-KODA | 12,13-epoxy-9-oxo-10(*E*)-octadecenoic acid |
| 225 | 59 | 2.44 | 12OH-JA | 12-hydroxy-jasmonic acid |
| 338.3 | 130.1 | 3.19 | 12OH-JA-Ile | 12-hydroxy-jasmonic acid isoluecine |
| 291.3 | 165 | 11.13 | 12-OPDA | 12-oxo-10(*Z*),15(*Z*)-phytodienoic acid |
| 295.3 | 195.2 | 13.46 | 13-HOD | 13(*S*)-hydroxy-9(*Z*),11(*E*)-octadecatrienoic acid |
| 293.2 | 113.2 | 14.16 | 13-KOD | 13-oxo-9(*Z*),11(*E*)-octadecadienoic acid |
| 309.4 | 180.8 | 10.32 | 13,12-KODA | 13-hydroxy-12-oxo-9(*Z*),15(*Z*)-octadecadienoic acid |
| 311.3 | 194.9 | 11.42 | 13,12-KOMA | 13-hydroxy-12-oxo-9(*Z*)-octadecenoic acid |
| 293.2 | 191.1 | 15.15 | 2-HOD | 2-hydroxy-9(*Z*),15(*Z*)-octadecadienoic acid |
| 327.1 | 137 | 5.57 | 9,10,11-THOD | 9(*S*),10(*S*),11(*R*)-trihydroxy-12(*Z*),15(*Z*)-octadecadienoic acid |
| 329.5 | 201 | 6.69 | 9,10,11-THOM | 9(*S*),10(*S*),11(*R*)-trihydroxy-12(*Z*)-octadecenoic acid |
| 327.1 | 137 | 4.94 | 9,10,13-THOD | 9(*S*),10(*S*),13(*S*)-trihydroxy-11(*E*),15(*Z*)-octadecadienoic acid |
| 329.5 | 139 | 5.8 | 9,10,13-THOM | 9(*S*),10(*S*),13(*S*)-trihydroxy-11(*E*)-octadecenoic acid |
| 313.2 | 201.2 | 10.46 | 9,10-DiHOM | threo-9,10-dihydroxy-12(*Z*)-octadecenoic acid |
| 326.9 | 229.1 | 5 | 9,12,13-THOD | 9(*S*),12(*S*),13(*S*)-trihydroxy-10(*E*),15(*Z*)-octadecadienoic acid |
| 329.5 | 210.9 | 5.74 | 9,12,13-THOM | 9(*S*),12(*S*),13(*S*)-trihydroxy-10(*E*)-octadecenoic acid |
| 295.3 | 171.1 | 13.53 | 9-HOD | 9(*S*)-hydroxy-10(*E*),12(*Z*)-octadecadienoic acid |
| 293.2 | 171.2 | 12.04 | 9-HOT | 9(*S*)-hydroxy-10(*E*),12(*Z*),15(*Z*)-octadecatrienoic acid |
| 293.2 | 195.1 | 12.25 | 9-KOD | 9(*S*)-oxo-10(*E*),12(*Z*)-octadecatrienoic acid |
| 291.3 | 184.7 | 13.04 | 9-KOT | 9(*S*)-oxo-10(*E*),12(*Z*),15(*Z*)-octadecatrienoic acid |
| 311.3 | 293.2 | 11.76 | 9,10-KOMA | 9-hydroxy-10-oxo-12(*Z*)-octadecadienoic acid |
| 309.1 | 291.3 | 9.02 | 9,12-KODA | 9-hydroxy-12-oxo-10(*E*),15(*Z*)-octadecadienoic acid |
| 311.3 | 293.1 | 10.29 | 9,12-KOMA | 9-hydroxy-12-oxo-10(*E*)-octadecenoic acid |
| 263.4 | 153.1 | 3.33 | ABA | Abscisic acid |
| 186.9 | 124.9 | 2.83 | AZA | Azelaic Acid |
| 121.1 | 76.9 | 2.54 | BA | Benzoic acid |
| 147 | 102.9 | 3.6 | CA | Cinnamic acid |
| 162.9 | 118.9 | 2.05 | COUMA | Coumaric acid |
| 263.1 | 165 | 8.29 | dn-12-OPDA | dinor-12-oxo-8(*Z*),13(*Z*)-phytodienoic acid |
| 174 | 129.9 | 2.77 | IAA | Indole-3-acetic acid |
| 209.1 | 59 | 4.24 | JA | (+)-7-iso-jasmonic acid |
| 322.2 | 130 | 6.82 | JA-Ile | (+)-7-Jasmonoyl-L-isoleucine |
| 237 | 165.1 | 6.38 | OPC-4 | 8-[3-oxo-2-cis-[(*Z*)-2-pentenylcyclopentyl]butanoic acid |
| 311.2 | 183.2 | 10.3 | 12,13-DiHOD* | threo-12,13-dihydroxy-9(*Z*),15(*Z*)-octadecenoic acid |
| 311.2 | 201.2 | 10.35 | 9,10-DiHOD* | threo-9,10-dihydroxy-12(*Z*),15(*Z*)-octadecenoic acid |
| 226.9 | 208.7 | 3.2 | 9OH-TAN* | 9-hydroxy-12-oxo-10(*E*)-dodecenoic acid |
| 223 | 195 | 7.3 | OTD* | 13-oxo-9(*Z*),11(*E*)-tridecadienoic acid |
| 137 | 92.9 | 2.45 | SA | Salicylic acid |
| 226.9 | 182.7 | 5.36 | TA | Traumatic acid |
| 210.9 | 182.7 | 6.56 | TAN | Traumatin |

[A-B] Select m/z ions for compound identification (see Methods section)
[C] Compound retention time

**Supplemental Figures**

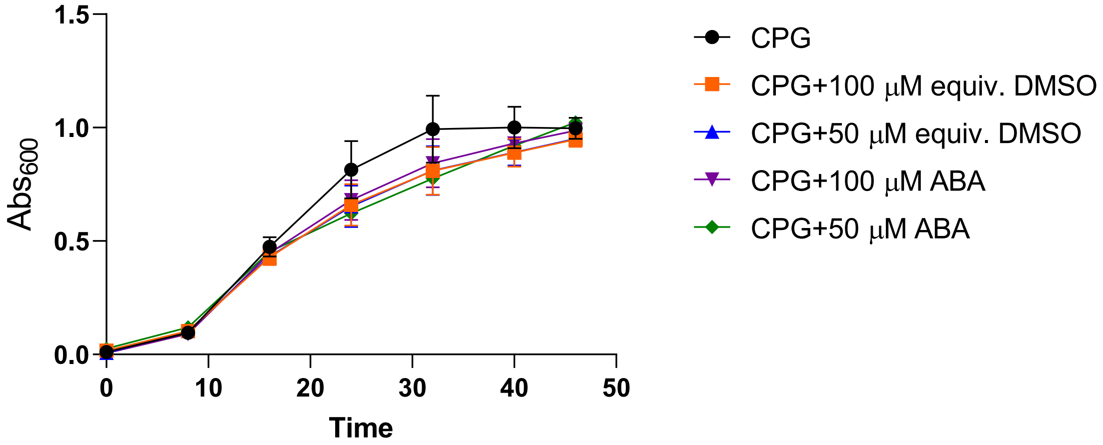


**Supplemental Figure 1. ABA and DMSO are not toxic to *Rs*.** Growth of *Rs* cultured in rich CPG broth supplemented with ABA in DMSO, or water with DMSO because ABA stock solutions were dissolved in DMSO (100 µM treatments contained 0.1 µL of either DMSO or ABA stock solution, and 50 µM treatments contained 0.05 µL DMSO or ABA stock solution) (ANOVA Area Under Curve/AUC, Fisher's LSD multiple comparisons to CPG control, 100 µM DMSO, *P*=.55; 50 µM DMSO, *P*=.53; 100 µM ABA, *P*=.67; 50 µM ABA, *P*=.45). Growth was measured spectrophotometrically using a Bio-Tek plate reader; the data represent eight technical reps/treatment. The bars represent the standard error of the mean.

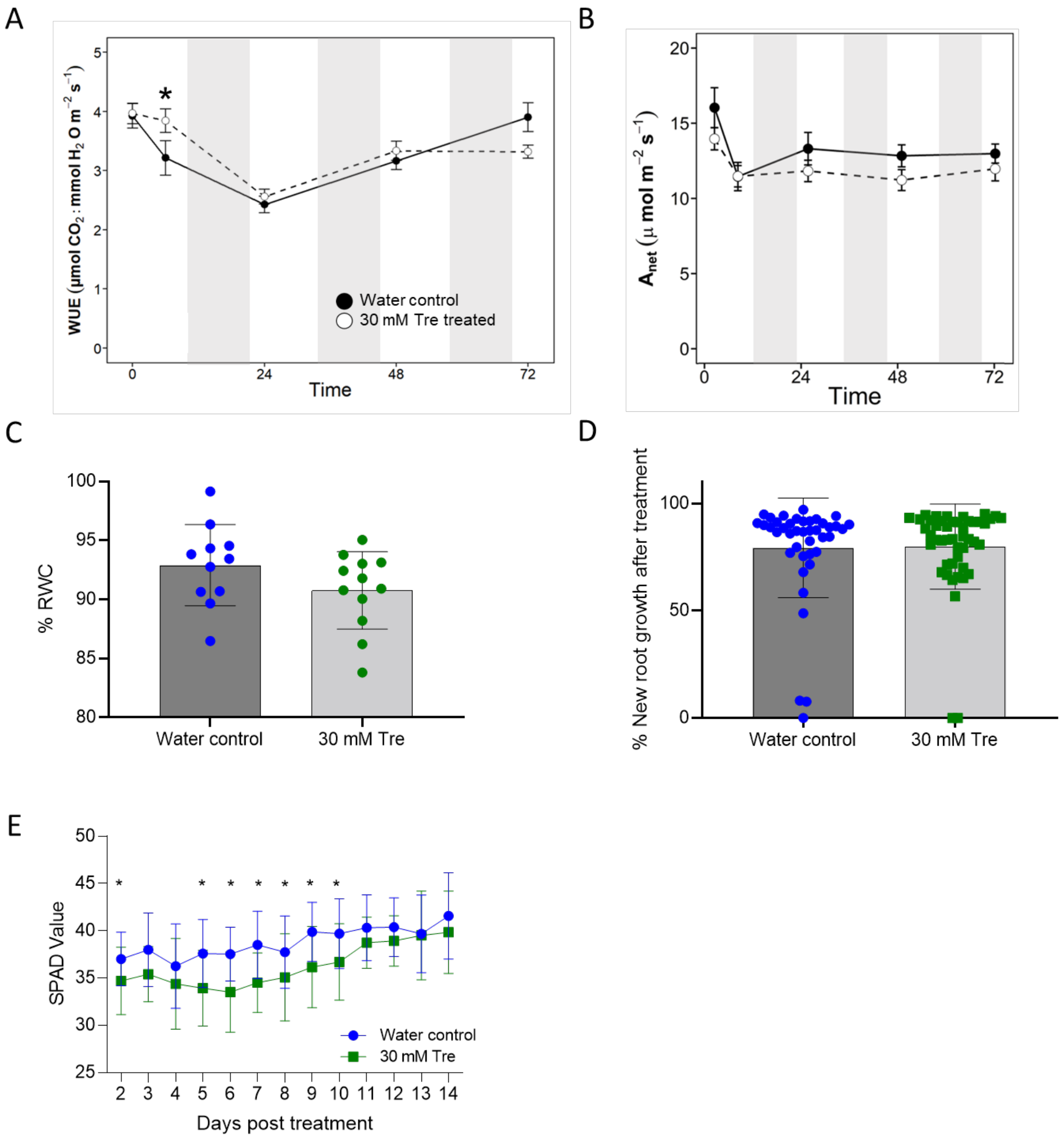


**Supplemental Figure 2. Trehalose treatment temporarily improves water efficiency without affecting photosynthesis; trehalose treatment does not affect root growth and relative water content of leaves but lowers chlorophyll concentration. A)** Water efficiency, measured with a LI-COR as photosynthetic rate/transpiration, in trehalose treated plants compared to water only controls (t-test, *P*<0.5). **B)** Photosynthesis, measured as $A_{net}$, in trehalose treated plants (Mixed model ANOVA, Tukey's HSD, *P*>.05). The water efficiency and photosynthetic data represent 20 plants/treatment. LI-COR measurements were taken ~10 min after treatment and then in the morning every time point after.

Shaded bars on the graphs represent night periods. All data were calculated from the same LICOR time points. The barson the data points represent the standard deviation. **C)** Relative water content of leaves was calculated from the fresh weight, hydrated weight, and dry weight of leaves from water control or trehalose treated BB tomatoes (t-test, $P$=.14). The data represent two biological replicates with five technical replicates (leaves)/treatment). The bars represent the standard deviation. **D)** ‘Bonny Best’ tomato seedlings were placed in root growth pouches, watered with 10mL of either water or trehalose for one day, then watered as needed for ten days. Root length was measured after ten days (t-test, $P$=.31). The data represent three biological replicates with ten roots/treatment. Percent of new root growth was calculated as %new root growth= (new growth/total root length)*100. The bars represent the standard deviation. **E)** SPAD meter measurements of chlorophyll content in leaves following trehalose treatment (2-way ANOVA Fishers LSD multiple comparison, * = $P$<0.5 or lower). The bars represent the standard deviation.

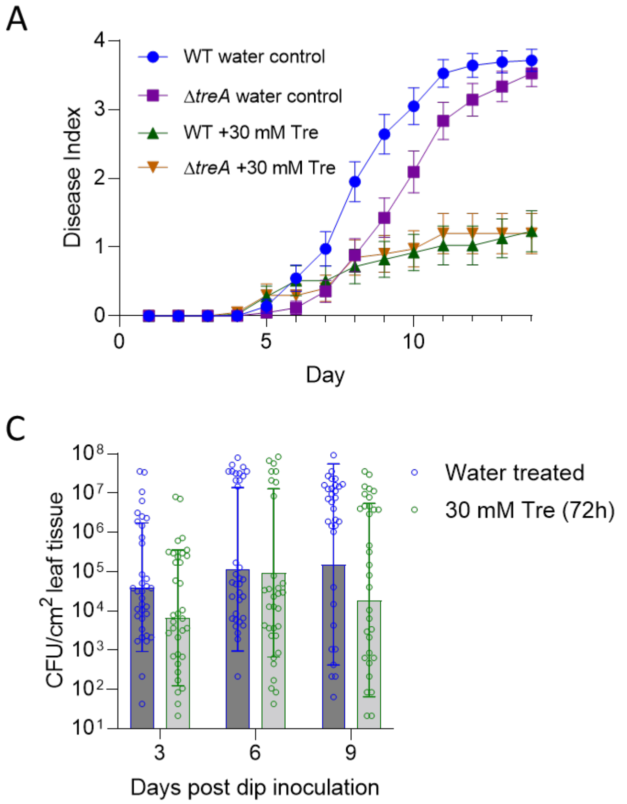


**Supplemental Figure 3. Trehalose treatment does not nutritionally enrich the soil environment for *R. solanacearum*, and its protection is limited to root application. A)** Disease progress curve of plants treated with trehalose or water, then infected with either wild-type *Rs* or a Δ*treA* mutant unable to catabolize trehalose (one-way ANOVA of areas under the curve, WT $H_2$0 vs. WT tre, *P*=.0008, Δ*treA* $H_2$0 vs. Δ*treA* tre, *P*=.017). The data represent three bioreps each containing 13-15 plants per treatment. The bars represent the standard error. **B)** Disease development in plants sprayed once with 30 mM trehalose or water, and then soil-soak inoculated with *Rs* 48 h later (ANOVA of AUC, Fisher's LSD multiple comparisons to $H_2$0 soil, $H_2$0 spray, *P*=.67; tre spray, *P*=.63; tre soil, *P*=.63). The data represent three biological replicates each containing fifteen plants per treatment. The bars represent the standard error. **C)** Colonization of tomato leaves by the bacterial leaf spot pathogen *Xanthomonas gardneri*; plants were dip-inoculated with *X. gardneri* 72 h after root treatment with water or 30 mM trehalose (Mann-Whitney, day 3, *P*=.06; day 6, *P*=.61; day 9, *P*=08). The data represent three biological replicates with twelve plants/treatment/timepoint. The graph displays the geometric means; bars indicate the geometric standard deviation of the data.

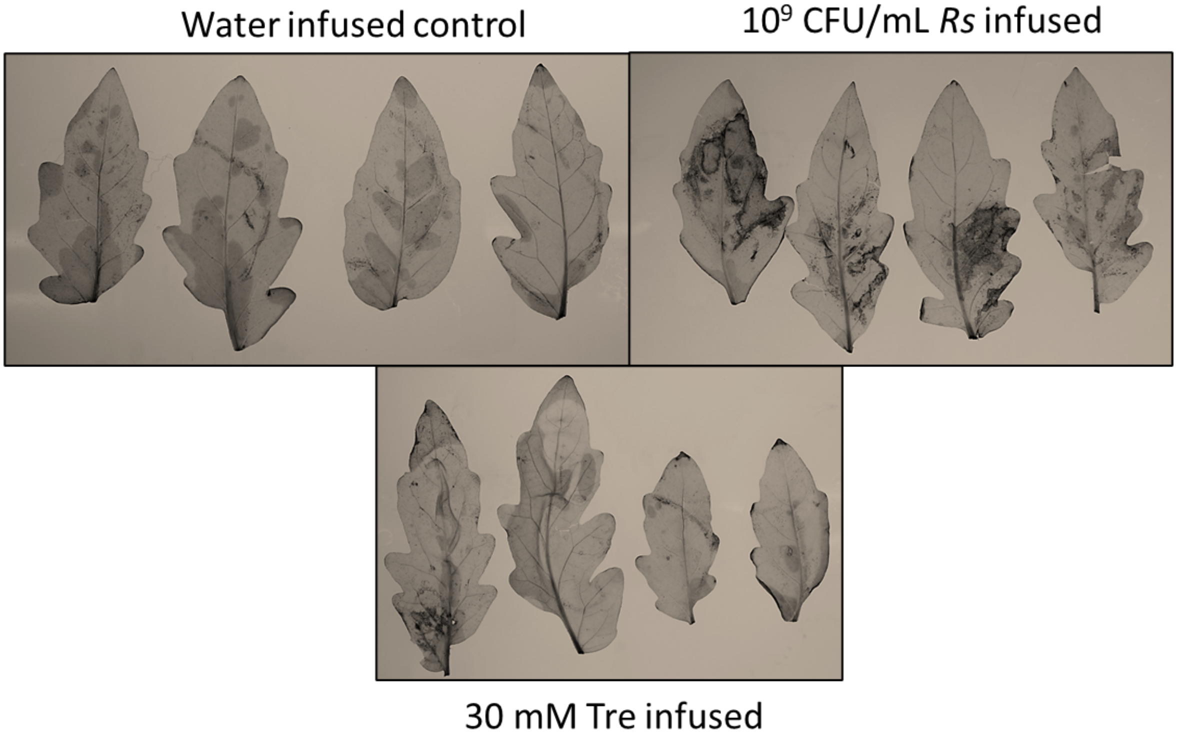


**Supplemental Figure 4. Infusing leaves with trehalose did not trigger ROS production in 'Bonny Best' tomato leaves.** DAB staining of 'Bonny best' tomato leaves infused with water, $10^9$ CFU/mL *Rs*, or 30 mM trehalose solution to assess the effect of trehalose treatment on ROS production. The data represent three biological replicates, with four plants per biological replicate per treatment. Photos are representative samples and images were uniformly sharpened 25% to increase contrast.
